# The SATB1-MIR22-GBA axis mediates glucocerebroside accumulation inducing a cellular senescence-like phenotype in dopaminergic neurons

**DOI:** 10.1101/2023.07.19.549710

**Authors:** Taylor Russo, Benjamin Kolisnyk, Aswathy Bs, Tae Wan Kim, Jacqueline Martin, Jonathan Plessis-Belair, Jason Ni, Jordan A. Pearson, Emily J. Park, Roger B. Sher, Lorenz Studer, Markus Riessland

## Abstract

Idiopathic Parkinson’s Disease (PD) is characterized by the loss of dopaminergic neurons in the substantia nigra pars compacta, which is associated with neuroinflammation and reactive gliosis. The underlying cause of PD and the concurrent neuroinflammation are not well understood. In this study, we utilized human and murine neuronal lines, stem cell–derived dopaminergic neurons, and mice to demonstrate that three previously identified genetic risk factors for PD, namely SATB1, MIR22HG, and GBA, are components of a single gene regulatory pathway. Our findings indicate that dysregulation of this pathway leads to the upregulation of glucocerebrosides (GluCer), which triggers a cellular senescence-like phenotype in dopaminergic neurons. Specifically, we discovered that downregulation of the transcriptional repressor SATB1 results in the derepression of the microRNA miR-22-3p, leading to decreased GBA expression and subsequent accumulation of GluCer. Furthermore, our results demonstrate that an increase in GluCer alone is sufficient to impair lysosomal and mitochondrial function, thereby inducing cellular senescence dependent on S100A9 and stress factors. Dysregulation of the SATB1-MIR22-GBA pathway, observed in both PD patients and normal aging, leads to lysosomal and mitochondrial dysfunction due to the GluCer accumulation, ultimately resulting in a cellular senescence-like phenotype in dopaminergic neurons. Therefore, our study highlights a novel pathway involving three genetic risk factors for PD and provides a potential mechanism for the senescence-induced neuroinflammation and reactive gliosis observed in both PD and normal aging.

## INTRODUCTION

The hallmark motor symptoms of Parkinson’s Disease (PD) arise primarily from the selective loss of dopaminergic (DA) neurons in the substantia nigra pars compacta of the midbrain, which are particularly vulnerable to degeneration (Dauer and Przedborski, 2003). Previously, our research identified the transcription factor Special AT-Rich Sequence-Binding Protein 1 (SATB1) as a genetic master regulator with a neuroprotective role specifically in nigral DA neurons (Brichta et al., 2015). Moreover, *SATB1* has been recognized as a genetic risk factor for PD (Hu et al., 2020; Nalls et al., 2019). In a recent study, we demonstrated that *SATB1* knockout (KO) triggers p21-dependent cellular senescence specifically in post-mitotic DA neurons (Riessland et al., 2019). Although the upregulation of p21 alone is capable of inducing senescence, inability of p21 knockdown to fully rescue the senescence phenotype in SATB1-KO cells indicates the involvement of an additional pathway.

Both human stem cell–derived SATB1-KO and mouse Satb1-knockdown DA neurons exhibited the classical hallmarks of senescence including mitochondrial damage and severe lysosomal dysfunction (Riessland, 2020; Riessland et al., 2019). Interestingly, we observed a significant downregulation of β-glucocerebrosidase (GCase), a critical lysosomal membrane protein encoded by *GBA*. This finding is of high interest as GBA is the most common genetic risk factor for PD (Sidransky et al., 2009). *GBA* encodes the enzyme GCase that cleaves the β-glucosidic linkage of glucosylceramides, also known as glucocerebrosides (GluCer) (Beutler, 1992). Decreased levels and activity of GCase have been observed in both normal aging and PD (Huh et al., 2021; Rocha et al., 2015). Notably, elevated GluCer levels in the cerebrospinal fluid (CSF) of idiopathic PD patients were associated with a rapid decline in Montreal Cognitive Assessment scores (Huh et al., 2021).

In this study, we investigated the SATB1-dependent regulation of GBA and the relationship between GluCer and cellular senescence, specifically in DA neurons. GCase ameliorated the SATB1-KO phenotype and elevated levels of GluCer alone were sufficient to induce a S100A9-mediated cellular senescence-like phenotype. In summary, we demonstrated that the loss of SATB1 led to the de-repression of microRNA (miRNA) miR22-3p, which in turn represses levels of GBA (Straniero et al., 2017). This novel pathway functionally connects SATB1, miR22-3p, and GBA. Furthermore, we demonstrated that the accumulation of GluCers alone leads to mitochondrial and lysosomal dysfunction, ultimately resulting in a cellular senescence-like state of DA neurons. These data suggest that the enzymatic activity of GCase is critical to DA neuron function.

## RESULTS

### Loss of SATB1 results in miR22-3p-mediated downregulation of GCase

Previously, we described the robust dysregulation of lysosomal gene expression in senescent SATB1-KO DA neurons (Riessland et al., 2019). Given the critical role of the lysosomal gene GBA in the molecular pathology of PD, we assessed GCase protein levels and observed a significant downregulation in SATB1-KO DA neurons compared to wild-type (WT) neurons (Figure 1a). Through chromatin immunoprecipitation sequencing (ChIP-seq) experiments, we found that the transcriptional repressor SATB1 binds with high abundance to the promoter region of *GBA* (Figure S1a), indicating that the removal of SATB1 could upregulate *GBA* expression. Consequently, at the RNA level, we observed a modest but significant upregulation of *GBA* following the elimination of SATB1 (Figure 1b), which appears contradictory to the significant downregulation of GCase at the protein level (Figure 1a).

**Figure 1.**
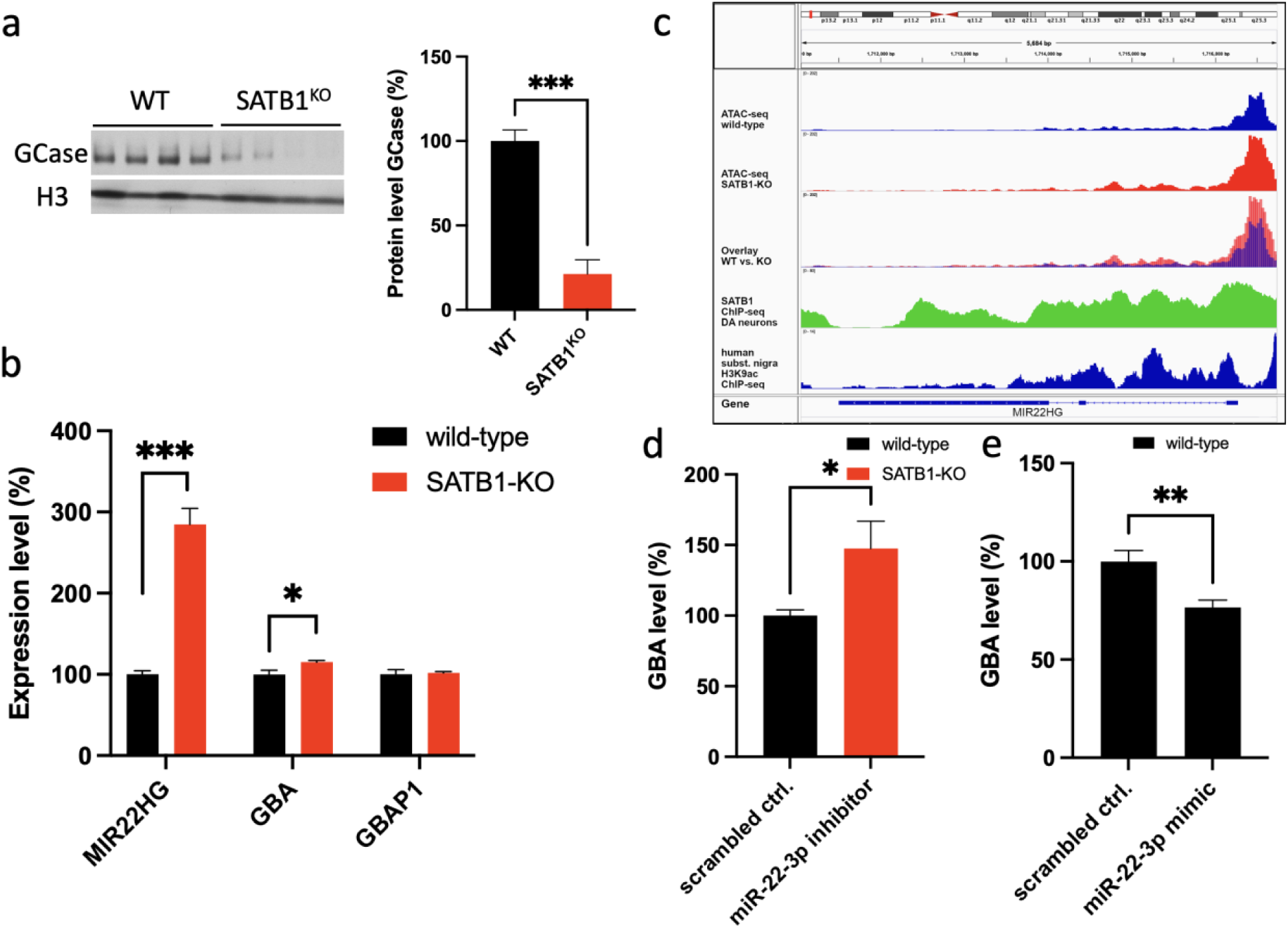
Knockout (KO) of SATB1 leads to MIR22HG de-repression and subsequent downregulation of GCase in dopaminergic (DA) neurons. **a**, Western blot analysis and quantification reveal a significant decrease in GCase protein levels in SATB1-KO compared to wild-type (WT) DA neurons (n=6). **b**, RNA expression profiling of MIR22HG, GBA, and GBAP1 in WT and SATB1-KO DA neurons shows differential expression (Day 50, n=3). **c**, Genomic data display the MIR22HG gene with ATAC-seq enrichment tracks from WT and SATB1-KO DA neurons, SATB1 ChIP-seq from WT DA neurons, and H3K9ac ChIP-seq data from human substantia nigra as a regulatory region marker. **d**, Quantification of GBA protein expression in SATB1-KO SK-N-MC cells treated with a miR22-3p inhibitor or scrambled control. **e**, Quantification of GBA protein expression in WT SK-N-MC cells treated with a miR22-3p mimic or scrambled control. Data are presented as mean ± S.E.M. Student’s t-tests were performed. * p<0.05; ** p<0.01; *** p<0.001.

To investigate this discrepancy, we performed ChIP-seq and RNA-seq analyses, to evaluate the potential of SATB1 to directly regulate known regulators of GBA. Doing so, we demonstrated the binding of SATB1 to the regulatory region of *MIR22HG* (miR22 host gene) (Figure 1c). This gene locus encompasses the GBA-regulatory miRNA miR22-3p (Straniero et al., 2017). The loss of SATB1 resulted in a 3-fold upregulation of miR22 expression (Figure 1b, c), providing support for a potential mechanism underlying the effects on GCase levels in SATB1-KO models. miRNAs can bind to RNA and inhibit its translation into protein. Thus, SATB1-KO leads to a mild increase in *GBA* RNA levels, a substantial increase in miR22-3p levels that sequester *GBA* RNA, and consequently, an overall decrease in GCase protein levels. Importantly, we ruled out the previously reported sponge effect mediated by *GBAP1*, as we did not observe any change in its expression (Figure 1b) (Straniero et al., 2017).

In addition to RNA-seq and SATB1-ChIP-seq analyses, we investigated chromatin accessibility using Assay for Transposase-Accessible Chromatin (ATAC)-seq data. The gene locus of *MIR22HG* exhibited significantly increased accessibility in SATB1-KO compared to WT DA neurons, suggesting reduced transcriptional repression (Figure 1c). To emphasize the tissue-specific importance of the regulatory gene region where SATB1 binds, we included H3K9ac ChIP-seq data derived from human substantia nigra (ENCODE, Figure 1c).

To validate our hypothesis that miR22-3p mediates the decrease in GCase upon SATB1 removal, we performed functional assays in the DA-like SK-N-MC human neuroblastoma cell line. The inhibition of mi22-3p upregulated GCase levels in SATB1-KO cells (Figure 1d), while treatment with a miR22-3p mimic downregulated GCase levels in WT cells (Figure 1e). Taken together, our data establish a functional pathway linking SATB1, miR22-3p, and GBA, which are all recognized genetic risk factors for PD.

### Overexpression of GBA ameliorates the vulnerability of SATB1-KO cells

Based upon our previous discovery of SATB1 as a neuroprotective genetic master regulator of DA neurons, we aimed to investigate whether the downregulation of GBA mediated by SATB1 increases cellular vulnerability. Due to their ease of transfection, we generated SATB1-KO Neuro-2A (N2A) murine cells. Using these cells, we confirmed that Satb1 regulates Gba levels (Figure S2a). As observed in the human system, the murine Satb1-KO cells exhibited a senescence-like phenotype, as indicated by an SA-b-Gal assay (Figure 2a, b), along with a significant decrease in GCase protein levels (Figure 2c). This reduction in GCase levels was consistent with a significant decrease in GCase enzymatic activity (Figure 2d). Furthermore, the vulnerability of Satb1-KO cells to both PD-specific and general toxins, such as 6-hydroxydopamine (6-OHDA) and H_2_O_2_, was heightened (Figure 2e, f). Importantly, the increased vulnerability observed after Satb1 removal was rescued by GBA overexpression (Figure 2g).

**Figure 2.**
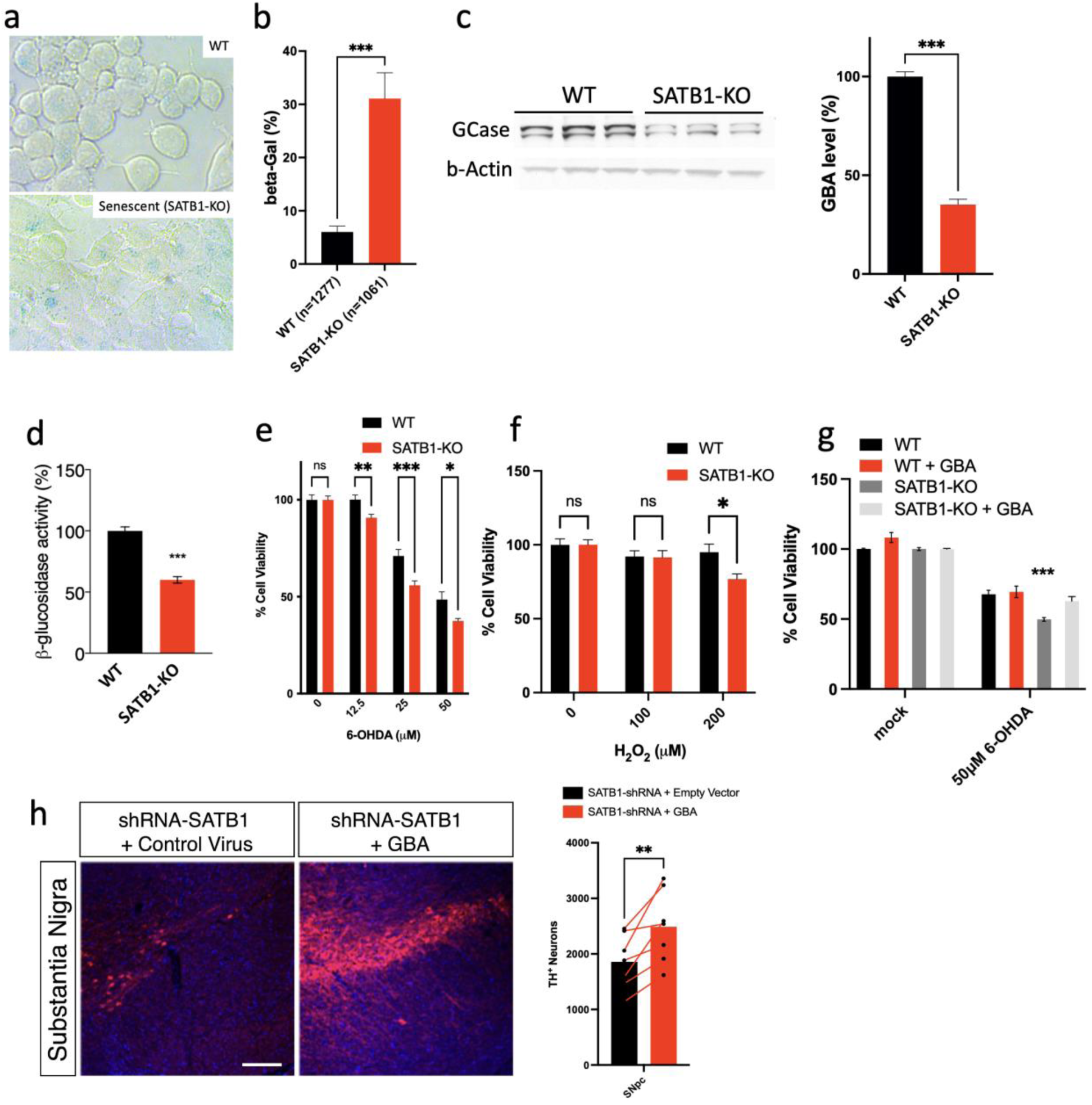
SATB1 knockout (KO) leads to reduced GCase levels and increased vulnerability *in vitro* and *in vivo.* **a,** Representative images from a SA-β-Gal assay comparing wild-type (WT) and SATB1-KO N2A cells. Blue cells indicate senescent cells. **b**, Quantification of the senescence assay in (**a**). WT, N=9 and n=1277; KO, N=10 and n=1061. **c**, Western blot analysis of GCase protein levels in N2A SATB1-KO cells confirms the reduction of GCase protein. **d**, Measurement of β-glucosidase enzymatic activity in N2A SATB1-KO cells compared to controls (n=24). Cell viability assays demonstrate the increased dose-dependent vulnerability of N2A SATB1-KO to treatment with 6-OHDA (**e**) and H_2_O_2_ (**f**) compared to control cells (n=24). **g**, Cell viability assays in WT and KO N2A cells with and without GBA overexpression. GBA overexpression rescues SATB1-KO vulnerability to 6-OHDA treatment. **h,** Representative tyrosine hydroxylase (TH) immunofluorescent staining and quantification in mice receiving a stereotaxic injection with a shRNA-Satb1 virus and a control vector, as well as a contralateral injection with a shRNA-Satb1 virus and a GBA-overexpressing virus. Injected *substantia nigra pars compacta* is shown and was quantified using unbiased stereological cell counting of TH^+^ cells (n=7) (scale bar: 500 um). Data are presented as mean ± S.E.M. Student’s t-tests were performed. ** p<0.01; *** p<0.001.

We have previously reported that the loss of Satb1 *in vivo* results in the loss of DA neurons (Riessland et al., 2019). To test the hypothesis that this loss of DA neurons was in part mediated by reduced GCase activity, we employed stereotactic co-injections of adeno-associated virus (AAV) of short-hairpin RNA (shRNA) targeting *Satb1* and an AAV expressing Gba in the midbrain of mice. Mice received a unilateral injection of either AAV-Satb1-shRNA and a control virus, or AAV-Satb1-shRNA and a virus encoding *Gba*. As previously observed, in the hemispheres treated with AAV-Satb1-shRNA and a control virus, a significant loss of tyrosine hydroxylase-positive DA neurons, hypothesized to be mediated by senescence, was observed, 3 weeks after viral injection. In contrast, in the hemisphere treated with AAV-Satb1-shRNA and a virus encoding *Gba*, the overexpression of GBA significantly mitigated this Satb1-knockdown-induced loss of DA neurons specifically in the substantia nigra (Figure 2h). These data corroborate the *in vitro* findings. Conversely, overexpression of GBA in the ventral tegmental area had no significant effects (Figure S2b), and no general effects were observed from virus injection or GBA overexpression (Figure S2c). Furthermore, virus expression *in vivo* was confirmed (Figure S2d).

Taken together, these results demonstrate that reduced GCase levels in SATB1-KO models contribute to the heightened vulnerability of the neuronal cell line, and the introduction of GBA effectively rescues this phenotype, both *in vitro* and *in vivo*. These findings provide additional evidence supporting the SATB1-GBA pathway and suggest its potential role in mediating vulnerability, senescence, and loss of DA neurons in PD.

### Lysosomal dysfunction in SATB1-KO DA neurons causes α-synuclein (α-SYN) accumulation and is rescued by GBA overexpression

We previously reported that human stem cell–derived SATB1-KO DA neurons undergo cellular senescence and exhibit lysosomal dysfunction (Riessland et al., 2019). To further investigate the lysosomal pathology in SATB1-KO DA neurons and its association with GBA levels, we conducted LysoTracker staining and a cathepsin-D assay. Our results revealed a significant accumulation of dysfunctional lysosomes in Satb1-KO cells (Figure 3a, b). Interestingly, the overexpression of either SATB1 or GBA rescued this phenotype, indicating that reduced GCase activity mediates the lysosomal dysfunction observed in SATB1-KO DA neurons.

**Figure 3.**
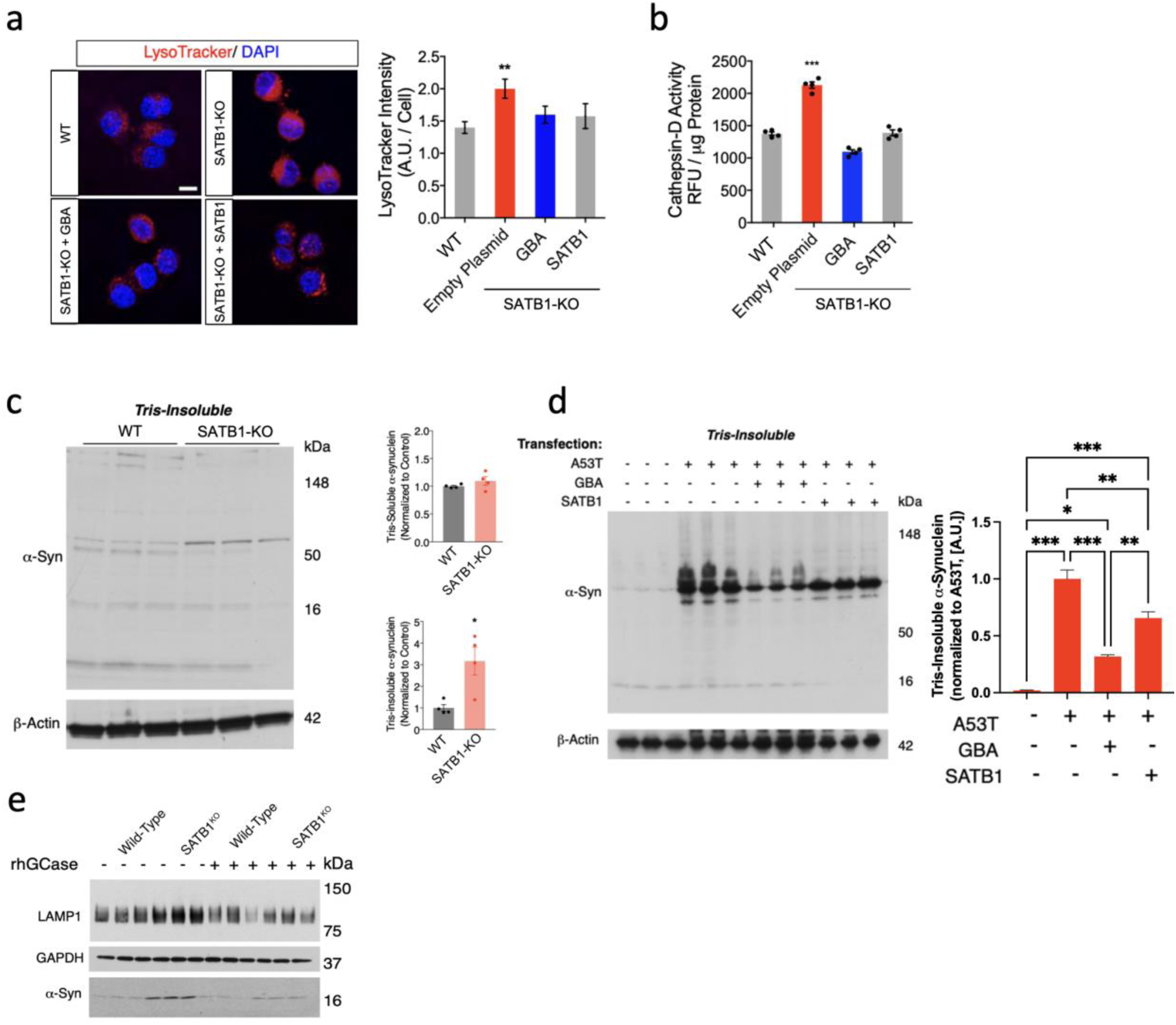
Knockout (KO) of SATB1 disrupts lysosomal function leading to α-SYN accumulation. **a,** Fluorescence microscopy images and quantification of lysosomal content in wild-type (WT) and SATB1-KO DA neurons with and without overexpression of GBA or SATB1. Lysosomal content (**a**) and cathepsin-D activity (**b**) are increased in SATB1-KO cells but can be normalized by extrinsic expression of either SATB1 or GBA. **c**, Triton X-100 insoluble α-SYN levels in N2A^WT^ and N2A^SATB1-KO^ cells. Representative Western blot and quantification are shown. **d**, Co-transfection of GBA or SATB1 along with α-SYN (A53T) reduced triton X-100 insoluble α-SYN when compared to A53T transfection alone. Representative Western blot and quantification are shown. **e,** Treatment of human DA neurons with recombinant GCase leads to a significant reduction in LAMP1 levels and normalizes the elevation of α-SYN (n=3). Data are presented as mean ± S.E.M. Student’s t-tests were performed. * p<0.05; ** p<0.01; *** p<0.001.

Reduced GCase function has been linked to glucosylsphingosine accumulation, in turn leading to the pathological aggregation of α-SYN (Taguchi et al., 2017). Therefore, we examined whether α-SYN aggregated in Satb1-KO neurons and found a significant elevation in the TRIS-insoluble form of α-SYN (Figure 3c). When the PD-mutant form of α-SYN (A53T) was overexpressed in WT cells, the TRIS-insoluble form also exhibited substantial aggregation. Notably, co-overexpression of either SATB1 or GBA significantly ameliorated this aggregation (Figure 3d), suggesting an improvement in the capacity of the lysosomal system to handle the mutant protein.

Considering the neuroprotective effect of GBA, we explored the therapeutic potential of recombinant GCase (rhGBA) by treating human stem cell–derived DA neurons lacking SATB1 with rhGBA. This treatment resulted in reduced accumulation of both lysosomal associated membrane protein 1 (LAMP1) and α-SYN in SATB1-KO DA neurons, indicating a beneficial effect on lysosomal function (Figure 3e). Overall, these findings support the presence of lysosomal impairment downstream of disruptions in the SATB1-GBA pathway, which can be rescued by the addition of GCase.

### Mitochondrial dysfunction and altered turnover in SATB1-KO DA neurons are rescued by GBA overexpression

In line with our previous report on SATB1-KO DA neurons entering cellular senescence with lysosomal dysfunction and mitochondrial impairment (Riessland et al., 2019), we investigated mitochondrial structure and function in the Satb1-KO N2A cell line. Transmission electron microscopy revealed structural abnormalities in mitochondrial cristae and membranes in Satb1-KO neuronal cells (Figure 4a, b). Measurements of oxygen consumption rate (OCR) and ATP production using a Seahorse XF Analyzer detected aberrations in mitochondrial function in Satb1-KO cells. Specifically, significant reductions in O_2_ respiration and ATP production rates were observed (Figure 4c). Mitotracker Red CMX analysis revealed a decrease in fluorescence in Satb1-KO cells, indicating decreased mitochondrial membrane potential (Figure 4d).

**Figure 4.**
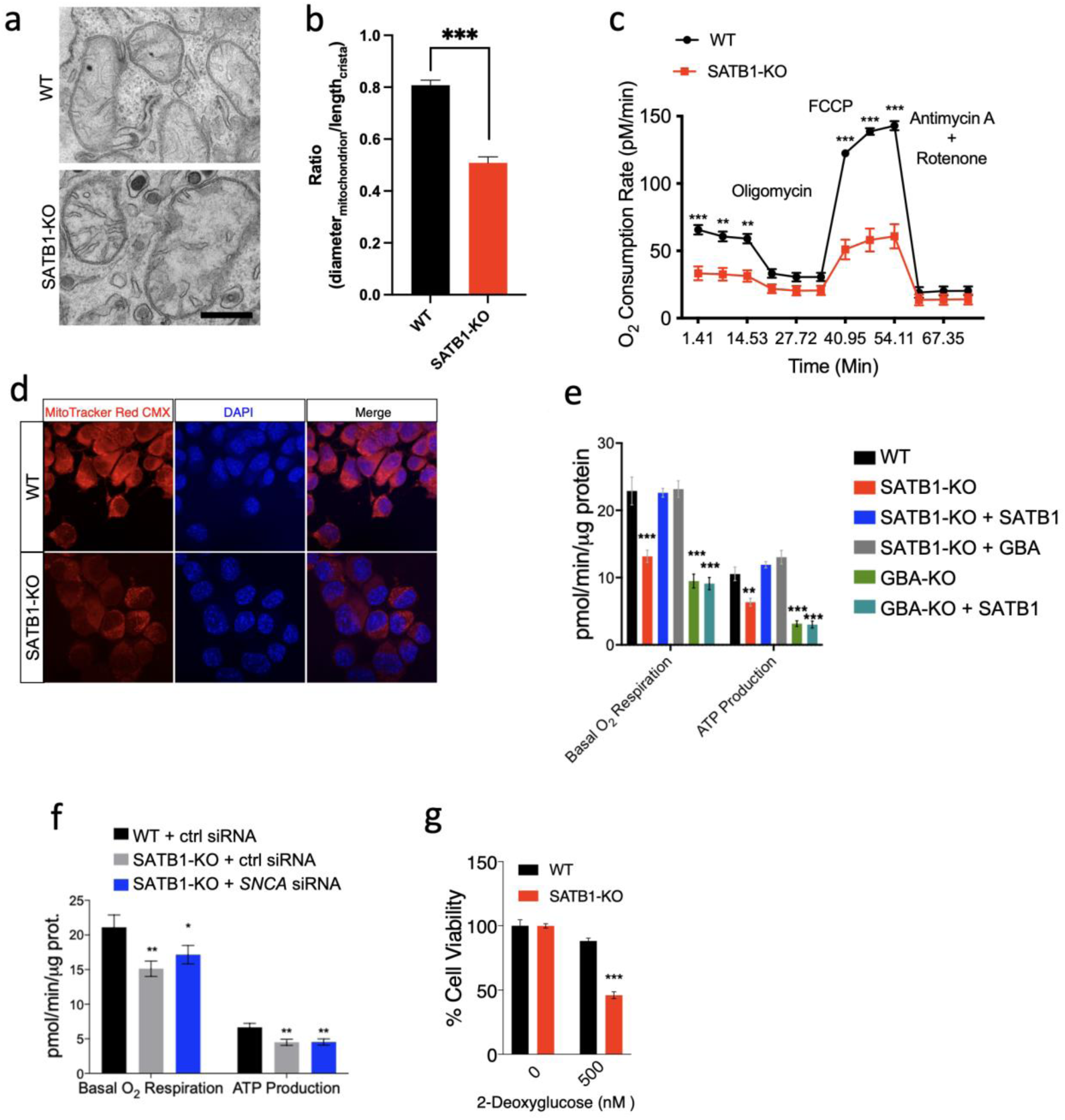
SATB1 knockout (KO) disrupts mitochondrial function and turnover. **a,** Transmission electron microscopy–based ultrastructural analysis of mitochondria in wild-type (WT) and SATB1-KO N2A cells. N2A^SATB1-KO^ cells exhibit severe accumulation of abnormal mitochondria. **b,** The ratio of mitochondrion diameter to cristae length was significantly lower in SATB1-KO DA neurons than in WT cells (WT, n=113; KO, n=106). **c,** Measurement of basal respiration and ATP production in N2A^WT^ cells transfected with control siRNA and N2A^SATB1-KO^ cells transfected with either control or α-SYN siRNA (n=12). **d,** Qualitative analysis of mitochondrial activity using a MitoTracker Red CMX probe in WT and N2A SATB1-KO DA neurons. **e,** Oxygen consumption rate measurement in N2A^WT^, N2A^SATB1-KO^, and N2A^GBA-KO^cells. SATB1 and GBA-KO cells show significant impairment in basal respiration and ATP production. Overexpression of SATB1 or GBA rescues ATP production and respiration, while SATB1 does not improve the phenotype in GBA-KO cells, indicating that SATB1 acts upstream of GBA. **f,** Downregulation of α-SYN is insufficient to restore oxygen respiration and ATP production in SATB1-KO DA neurons. **g,** Increased vulnerability of N2A^SATB1-KO^ cells to the glycolysis inhibitor 2-deoxyglucose compared to controls (n=6). Data are presented as mean ± S.E.M. Student’s t-tests were performed. * p<0.05; ** p<0.01; *** p<0.001.

Importantly, overexpression of GBA or SATB1 rescued the decreased O_2_ respiration and ATP production rates in Satb1-KO cells, while overexpression of SATB1 in Gba-KO cells had no effect (Figure 4e). Downregulation of α-SYN did not restore O_2_ respiration and ATP production rates in Satb1-KO cells, suggesting an α-SYN-independent mechanism (Figure 4f). Western blot analysis confirmed the knockdown of α-SYN (Figure S4f). Interestingly, native gel electrophoresis revealed a loss of integrity in oxidative phosphorylation complexes in both Satb1-KO and Gba-KO cells (Figure S4g, h). These findings further support the involvement of GBA in mediating structural and functional abnormalities of mitochondria in SATB1-KO models.

Mitochondrial quality control and mitophagy are critical pathways in PD, connecting lysosomes and mitochondria (Chung et al., 2016; Kamienieva et al., 2021). The interaction between phosphatase and tensin homolog (PTEN)-induced kinase 1 (PINK1) and parkin is essential for mitochondrial quality control. PINK1 binds to the surface of mitochondria, assessing membrane potential and activating the ubiquitin ligase parkin, which labels the mitochondrion for degradation by mitophagy (Eiyama and Okamoto, 2015). Immunostaining for parkin revealed a significant increase in parkin-positive mitochondria in Satb1-KO cells, indicating the accumulation of dysfunctional mitochondria (Figure S4a, b). Treatment with the mitochondrial uncoupler CCCP increased Pink1 levels in WT cells, and this effect was reversible upon washout of CCCP. In contrast, Satb1-KO cells exhibited a constant elevation of Pink1 that remained unaltered by CCCP treatment (Figure S4c). To investigate whether Satb1-KO cells, which lack functional mitochondria, rely on glycolysis, we subjected them to a 2-deoxyglucose treatment, which is toxic for glycolytic cells. As expected, we observed a significant increase in cell death in the Satb1-KO cells (Figure 4g).

Lastly, mitochondrial turnover was examined using the pH-dependent mitochondria-specific probe mKeima (Katayama et al., 2011). Intact mitochondria in the cytosol were labeled green, while damaged mitochondria in lysosomes were labeled red. In WT cells, treatment with the cellular stressors rotenone or 6-OHDA caused mitochondrial damage and translocation into lysosomes, leading to an increase in the mitophagy index. However, in Satb1-KO cells, there was no change in the mitophagy index (Figure S4d, e). These results support our hypothesis that mitochondrial impairment is a downstream effect of the disruption in the SATB1-GBA pathway. Given that both mitochondrial and lysosomal dysfunction are hallmarks of senescent cells, particularly senescent DA neurons in PD (Riessland et al., 2019), our findings suggest that the reduction in GCase may contribute to the senescence-like phenotype of SATB1-KO DA neurons through its induction of mitochondrial and lysosomal impairment.

### GluCer accumulates in SATB1-KO DA neurons

Next, we conducted transmission electron microscopy to characterize SATB1-KO DA neurons and observed a significant accumulation of intracellular lipid vesicles (Figure 5a, b). Given that GCase is responsible for cleaving the β-glucosidic linkage of GluCer (Beutler, 1992), we hypothesized that these lipid vesicles represent an accumulation of GluCer. To confirm our hypothesis, we performed immunocytochemistry and dot blot analyses using a GluCer-specific antibody in human stem cell–derived DA neurons. Compared to wild-type controls, there was a significant increase in GluCer levels in SATB1-KO DA neurons (Figure 5c, d). These results were confirmed in the murine Satb1-KO N2A cells (Figure S5a).

**Figure 5.**
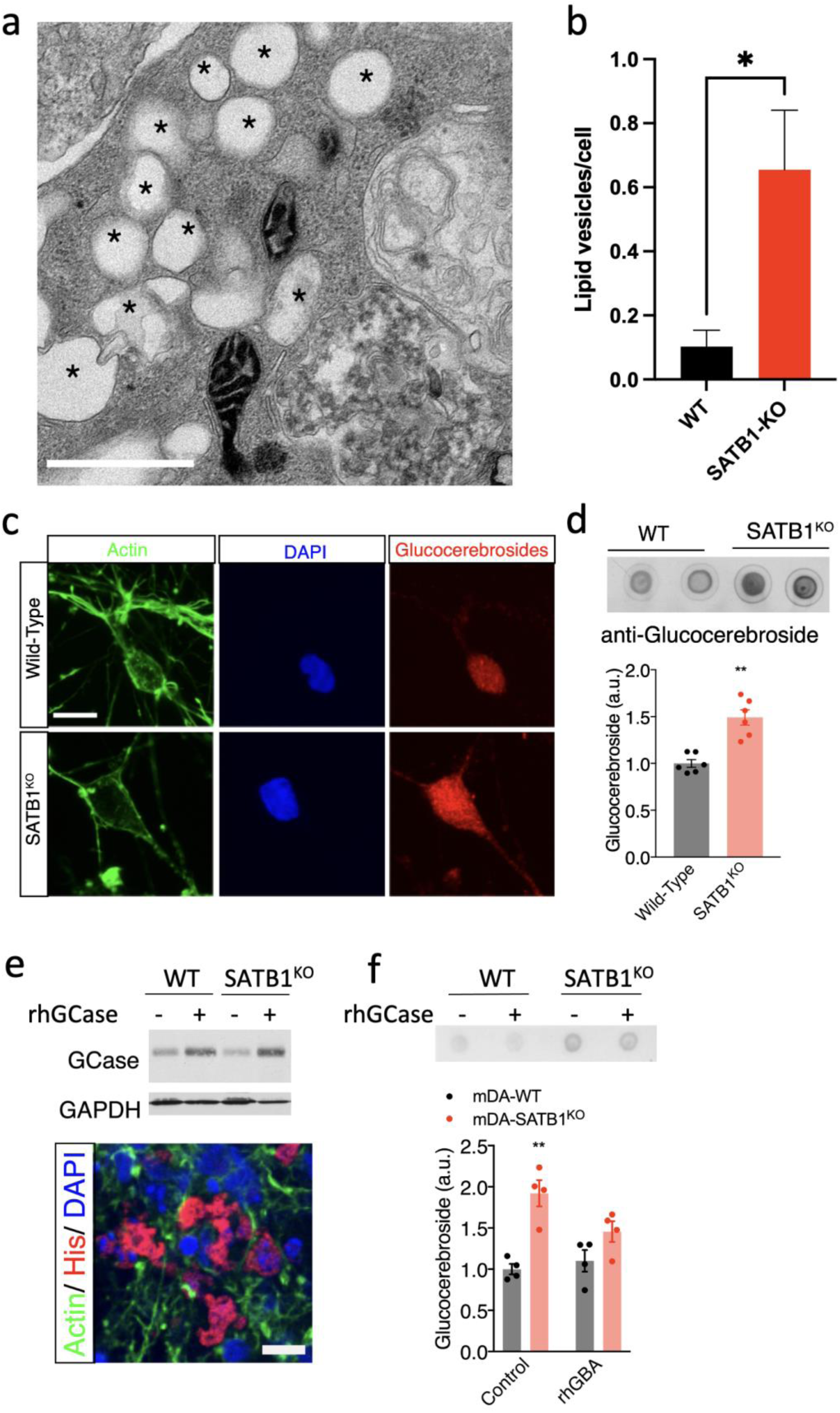
SATB1-KO-mediated GBA reduction causes lipid accumulation in DA neurons. **a**, Transmission electron microscopy (TEM) image showing lipid accumulation in SATB1^KO^ DA neurons (* indicates lipid inclusion). **b**, SATB1-KO DA neurons show a significant increase in the number of lipid vesicles per cell (cells analyzed for TEM: WT, n=88; KO, n=110). **c**, Immunofluorescence images using anti-GluCer antibodies and ActinGreen demonstrating increased GluCer levels in SATB1-KO DA neurons. **d,** Dot blot analysis and quantification of GluCer levels in SATB1-KO DA neurons relative to controls (normalized to protein amount, n=4). **e,** Treatment of human DA neurons with recombinant GCase leads to the presence of intracellular recombinant GCase. **f,** Significant reduction of GluCer accumulation in human DA neurons following treatment with recombinant GCase (n=4). Data are presented as mean ± S.E.M. Student’s t-tests were performed. * p<0.05; ** p<0.01.

To further investigate the consequence of GluCer accumulation, we also treated human stem cell–derived DA neurons with various concentrations of GluCer and performed dot blot assays to compare GluCer accumulation levels with those in SATB1-KO DA neurons. Treatment of WT DA neurons with 2.5 or 40 uM GluCer led to a significant increase in GluCer accumulation, which was comparable to that in SATB1-KO DA neurons (Figure S5b). Additionally, treatment of SATB1-KO DA neurons with rhGBA significantly reduced GluCer accumulation (Figure 5e, f). Moreover, we observed a non-significant but trending increase in GluCer following miR22-3p inhibition and a decrease in GluCer following treatment with a miR22-3p mimic in SK-N-MC cells (Figure S5c); these results are in line with the effects of miR22-3p on GCase levels (Figure 1d, e). Taken together, these data support our hypothesis that GluCer accumulation is a downstream effect of SATB1-KO, reinforcing the role of the SATB1-GBA pathway and the association between decreased GCase levels and GluCer dysregulation.

### GluCer directly induces lysosomal and mitochondrial impairment and triggers a cellular senescence-like phenotype

Having observed a significant accumulation of GluCer, our focus shifted to investigating the direct impact of GluCer elevation on lysosomal and mitochondrial dysfunction and its potential role in inducing a cellular senescence phenotype in DA neurons. Senescent cells are characterized by lipid accumulation, particularly ceramides (Venable et al., 2006), and the introduction of exogenous ceramides has been shown to induce senescence (Hannun and Obeid, 2008; Venable et al., 1995). Therefore, we aimed to explore whether accumulated GluCer in DA neurons directly contributes to lysosomal and mitochondrial impairment, bridging the SATB1-GBA pathway with GluCer dysregulation, mitochondrial and lysosomal dysfunction, and cellular senescence.

Treatment of WT DA neurons with 2.5 or 40 uM GluCer resulted in a significant increase in GluCer accumulation, statistically comparable to that observed in SATB1-KO DA neurons (Figure S5b). Lysotracker analysis in WT N2A cells following a 6-day treatment with 2.5 or 40 uM GluCer revealed a significant accumulation of lysosomes (Figure 6a), mirroring the phenotype of Satb1-KO cells (Figure 3a). To further elucidate the impact of GluCer treatment on mitochondrial function, human stem cell–derived DA neurons were treated with increasing concentrations of GluCer and subjected to an oxidative phosphorylation assay using a Seahorse XF Analyzer–based approach. We observed a dose-dependent impairment of mitochondrial function correlated with increasing GluCer concentrations (Figure 6b). Enzymatic inhibition of GluCer synthesis using GENZ-112638 alleviated this effect (Figure 6c). Collectively, these experiments provide evidence that GluCer accumulation directly impairs mitochondrial and lysosomal function in DA neurons.

**Figure 6.**
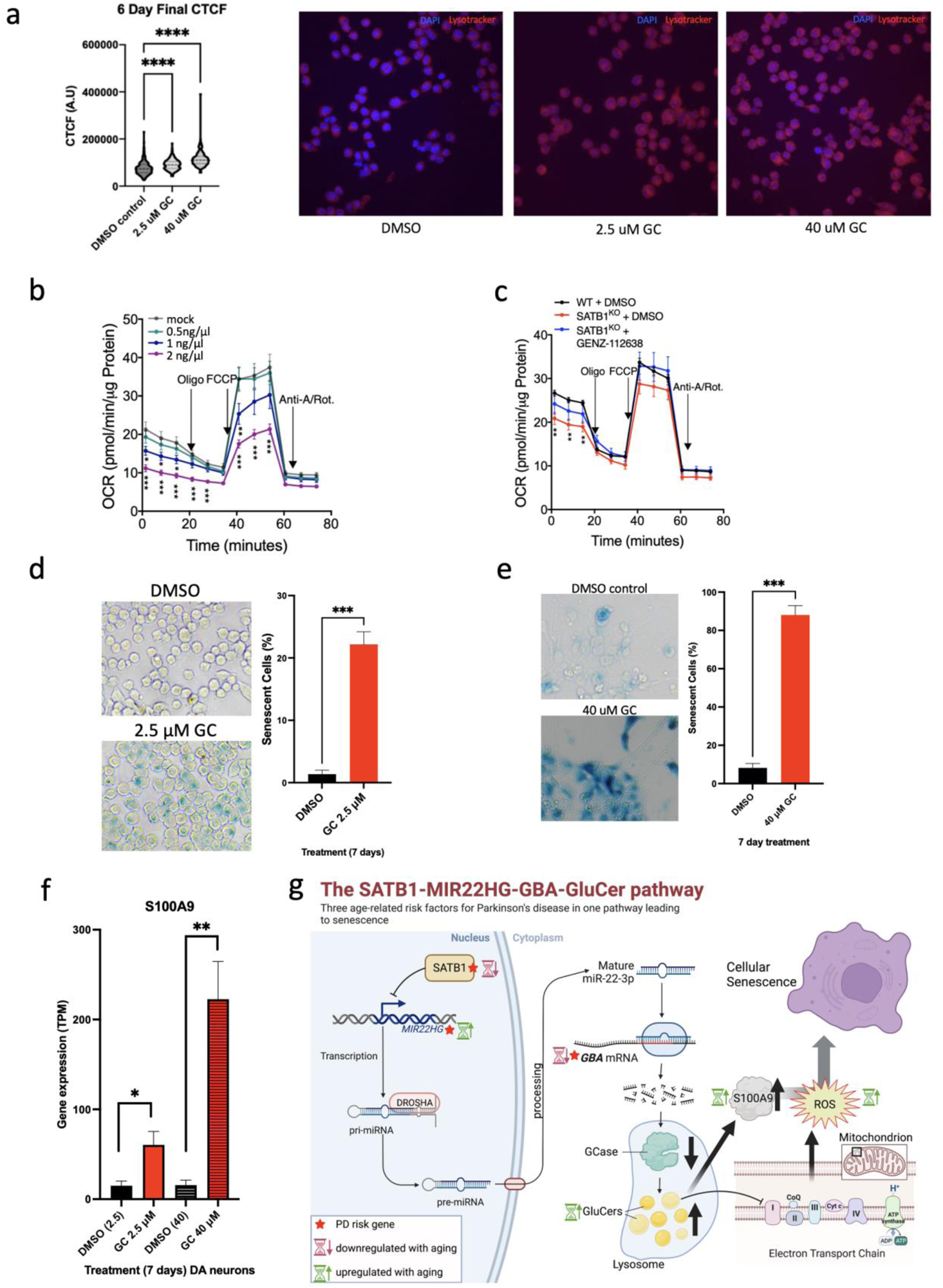
GluCer directly impairs mitochondria and lysosome function leading to S100A9 upregulation and a senescence-like phenotype in dopaminergic (DA) neurons. **a,** LysoTracker staining of wild-type (WT) N2A cells treated with DMSO control, 2.5 mM GluCer, or 40 mM GluCer for 6 days, demonstrating a significant dose-dependent increase in lysosomal content (corrected total cellular fluorescence) upon lipid treatment. **b,** Seahorse analysis of lipid-treated mesencephalic DA (mDA) neurons showing dose-dependent alterations in oxygen consumption rate (2 ng/ml is 2.8 mM). **c,** Treatment with GENZ-112638, an inhibitor of GluCer synthesis, mitigates the effects of lipid treatment on mitochondrial function. **d, e** SA-b-Gal-based senescence assay performed to evaluate senescence induction after 7 days of GluCer treatment in N2A cells (**d**) and mature WT human mDA neurons (**e**), demonstrating that GluCer treatment is sufficient to induce senescence in both cell types. **f,** RNA-seq analysis of WT human DA neurons treated with DMSO, 2.5 mM GluCer, or 40 mM GluCer, revealing increased S100A9 expression following a 7-day lipid treatment **g, Overview of the SATB1-MIR22HG-GBA-GluCer pathway.** The senescence regulator SATB1 acts as a negative regulator of MIR22HG expression. Decreased SATB1 levels lead to increased MIR22HG expression. Following nuclear processing, miR-22-3p targets GBA mRNA, reducing its transcription. This results in reduced GCase activity and the consequent accumulation of its substrate GluCer, a cerebroside that impairs mitochondrial and lysosomal function. GluCer accumulation leads to elevated ROS production and increased S100A9 expression, ultimately driving cellular senescence. Importantly, the genes SATB1, MIR22HG, and GBA (red stars) are associated with Parkinson’s Disease. During aging, SATB1 and GBA levels decrease (red hourglasses), while MIR22HG, GluCer, S100A9, and ROS levels increase (green hourglasses), rendering DA neurons more vulnerable. Both S100A9 and miR22 induce cellular senescence (Shi et al., 2019; Xu et al., 2011).

Given our previous findings on the involvement of SATB1 in cellular senescence in DA neurons, our next objective was to investigate whether elevated GluCer levels alone could induce a similar phenotype. As expected, treatment of human stem cell–derived DA neurons and N2A cells with increasing concentrations of GluCer resulted in a significant increase in the number of senescent-like cells (Figure 6d, e). When mature WT human DA neurons were exposed to 2.5 or 40 uM GluCer, we observed a significant and dose-dependent increase in S100A9 expression compared to DMSO-treated control cells (Figure 6f). This is of particular interest since S100A9 is implicated in the induction of cellular senescence (Shi et al., 2019) and is significantly upregulated in multiple aging murine and human tissues, including those in the central nervous system (Swindell et al., 2013). Thus, this finding provides a potential explanation for the partial rescue of p21-dependent senescence observed in SATB1-KO DA neurons upon p21 inhibition (Riessland et al., 2019), suggesting that the upregulation of S100A9 induced by GluCer may directly drive a senescence-like phenotype in these cells, as we did not observe any significant upregulation of the expression of p21 (Figure S5d). These results establish a direct link between GluCer accumulation, lysosomal impairment, mitochondrial dysfunction, S100A9 upregulation, and cellular senescence, all of which occur downstream of the SATB1-GBA pathway. Our findings shed light on senescence as an age-related mechanism contributing to the dysfunction and heightened vulnerability of DA neurons.

## DISCUSSION

In this study, we present evidence that highlights the regulatory relationship of one genetic risk factor for PD by another: GBA by SATB1 (Nalls et al., 2019; Sidransky et al., 2009). This regulation is mediated through the direct control of the miRNA gene *MIR22HG* by SATB1, which itself is significantly influenced by the PD risk variant rs1109303 (Liu et al., 2019). Notably, the mature miRNA miR-22-3p plays a role in downregulating the expression of GCase, the protein product of *GBA*. Considering that PD is an age-related disorder, it is noteworthy that both GBA and SATB1 expression decline with age, while miR-22-3p expression increases (Ravanidis et al., 2020). Furthermore, significantly elevated levels of miR-22-3p have been observed in the plasma of individuals with REM sleep behavior disorder, a highly predictive condition for PD, as well as in those with Lewy body disease (Soto et al., 2022).

Relevant to the aging process and PD risk, GCase activity progressively decreases in the substantia nigra and putamen during normal aging. In sporadic (non-GBA mutant) PD patients who are in their sixth decade of life, GCase activity is reduced by approximately 50%, comparable to control individuals in their eighties (Rocha et al., 2015), suggesting a potential contribution to age-related PD risk. Additionally, glycolipid levels are elevated in the substantia nigra of these patients (Rocha et al., 2015). Collectively, these findings underscore the intricate interplay between GBA, SATB1, miR-22-3p, aging, and PD. The regulatory mechanisms and alterations in glycolipid metabolism may contribute to age-related susceptibility to PD, providing valuable insights into the pathogenesis of the disease.

In this study, we demonstrate that the accumulation of GluCer, the substrate of GCase, directly impacts the function of lysosomes and mitochondria in DA neurons, leading to the induction of S100A9 expression and a cellular senescence-like phenotype. S100A9, a senescence inducer, is upregulated with age (Swindell et al., 2013). It also co-localizes and co-aggregates with α-SYN in 20% of Lewy bodies and 77% of neuronal cells in the substantia nigra of PD patients (Horvath et al., 2018). Additionally, GBA-KO cells that exhibit elevated GluCer levels demonstrate upregulation of S100A9 (Gehrlein et al., 2023).

Ultrastructure and mitochondrial functional analyses in DA neurons revealed the colocalization of accumulating lipids with mitochondria, impairing oxidative phosphorylation. We observed dose-dependent impairments in basal respiration, ATP production, and maximal respiration due to GluCer accumulation, ultimately leading to a cellular senescence phenotype. This finding aligns with a recent report showing that specific knockout of Ndufs2 in DA neurons disrupts mitochondrial complex I and induces progressive parkinsonism, indicating the onset of cellular senescence (González-Rodríguez et al., 2021). Importantly, this study demonstrated that impairment of complex I resulted in a progressive loss of the DA phenotype, primarily observed in nigrostriatal axons, while promoting neuronal survival, indicating the initiation of cellular senescence. Furthermore, our Seahorse analyses of the oxidative phosphorylation of GluCer-treated DA neurons (Figure 6b) yielded findings consistent with previous studies on NDUFS2 KO cells, which also exhibit complex I impairment (Bandara et al., 2021). These results suggest that GluCer accumulation inhibits the function of complex I.

Mitochondrial dysfunction and elevated reactive oxygen species (ROS) production are well-documented characteristics of senescent cells (Martini and Passos, 2023). The importance of mitochondrial function in DA neurons is highlighted by studies that connect mutations in genes responsible for maintaining mitochondrial health to early-onset or autosomal dominant forms of PD (Beilina and Cookson, 2016; Haelterman et al., 2014). Environmental toxins and α-SYN, a major component of Lewy pathology, also inhibit complex I, thereby increasing PD risk (Faustini et al., 2017).

Our results show that lysosomal impairment is a downstream effect of the newly identified SATB1-MIR22-GBA pathway. We show the accumulation of dysfunctional lysosomes in a SATB1-KO model, which can be rescued by overexpressing SATB1 or GBA (Figure 3a, b). Additionally, we observed elevated levels of TRIS-insoluble α-SYN in Satb1-KO cells, indicating further impairment in lysosomal function. We successfully alleviated α-SYN accumulation in DA neurons through recombinant GCase treatment (Figure 3c, e). Recent findings suggest that reduced GCase activity leads to senescence in lysosomal disorders (Squillaro et al., 2017).

Overall, our findings demonstrate that accumulation of GluCer triggers a lysosomal and mitochondrial dysfunction-dependent senescence-like phenotype, indicating that cellular senescence may serve as a common mechanism underlying various PD insults. The presence of the senescence-associated secretory phenotype (SASP) in senescent cells, known to elicit immune reactions, could potentially explain the reported prodromal inflammation in the midbrain of incipient PD patients and suggest that killer T-cells may be involved in the loss of DA neurons (Galiano-Landeira et al., 2020; Sulzer et al., 2017).

In summary, our study demonstrates the dysregulation of the SATB1 → MIR22HG → GBA → GluCer → S100A9 → senescence pathway, which plays a critical role in aging DA neurons and the associated risk of PD development (Figure 6g). This novel insight paves the way for potential therapeutic approaches targeting senescence, such as the utilization of senolytics, to mitigate the risk and progression of PD. Further exploration of the connection between PD pathology, inflammaging, and cellular senescence is warranted to deepen our understanding of the disease and identify additional therapeutic opportunities.

## ACKNOWLEDGMENTS

This work was supported in part through grants 1R01NS124735-01A1 (M.R.) and States Army Medical Research and Material Command (USAMRMC) under Award No. W81XWH-12-1-0039 (M.R.), and by Aligning Science Across Parkinson’s [Grant number: ASAP-000472] and by core grant P30 CA08748. T.W.K. was supported in part by a Druckenmiller fellowship from the New York Stem Cell Foundation. Mitochondrial respiration measurements were performed using the Agilent Seahorse XFe96 analyzer with generous support from the Rockefeller University High-Throughput Screening and Spectroscopy Resource Center (HTSRC). We thank the Genomics Resource Center and the Electron Microscopy Resource Center (EMRC) at Rockefeller University and the Neurobiology and Behavior Imaging Core at Stony Brook University for technical support. Alphasynuclein-A53T was a gift from David Rubinsztein (Addgene plasmid # 40823; http://n2t.net/addgene:40823; RRID:Addgene_40823) (Furlong et al., 2000). pHAGE-mt-mKeima was a gift from Richard Youle (Addgene plasmid # 131626; http://n2t.net/addgene:131626; RRID:Addgene_131626) (Vargas et al., 2019). Opinions, interpretations, conclusions, and recommendations are those of the author and are not necessarily endorsed by the sponsors.

## MATERIALS AND METHODS

### Western blotting

Western blotting was performed as previously described (Riessland et al., 2019). In short, protein lysates were derived from cell cultures using RIPA buffer (#89900, Thermo Scientific) containing both protease and phosphatase inhibitors (#11836170001, Roche). To determine the protein concentration of each sample, a BCA assay (Thermo Scientific) was performed using a SpectraMax iD3 (Molecular Devices) plate reader. Equal quantities of protein were boiled in NuPAGE LDS sample buffer (Invitrogen) or Tris-Glycine-SDS sample buffer (Novex) at 95°C for 5 min and separated using 4–20% Tris-glycine (ThermoFisher, #EC6025 or Invitrogen # XP04200). Proteins were transferred to nitrocellulose membranes (BioRad) using a wet blotting method, blots were then blocked with 5% BSA (Jackson immuno) for 1 hour at room temperature, and subsequently incubated with the respective primary antibody overnight at 4°C. The following primary antibodies were used: rabbit monoclonal LAMP1 (CST #D2D11, 1:3000), rabbit monoclonal anti-SATB1 antibody (Abcam, # ab189847; 1:3000), rabbit polyclonal anti-GBA (Abcam #ab128879, #ab70004, 1:3000) or mouse polyclonal anti-GBA (Abnova #146-235, 1:1,000), rabbit monoclonal anti-beta Actin (CST #D6A8, 1:15000), rabbit polyclonal anti-alpha synuclein (CST #2642, 1:3000), rabbit polyclonal anti-PINK1 (Abcam #ab23707; 1:3000), rabbit polyclonal anti-GAPDH (Abcam # ab9485, 1:3000), rabbit polyclonal anti-Histone H3 (CST #D1H2, 1:5,000), and rabbit polyclonal anti-glycosyslceramide (Glycobiotech #RAS_0010, 1:3,000). Primary antibody was detected with HRP-linked donkey anti–rabbit IgG (GE Healthcare, #NA934V; 1:10,000) or HRP-liked sheep anti-mouse IgG (GE Healthcare, #NA931V, 1:10,000) together with Western Lightning Plus-ECL (Perkin Elmer, #NEL105001EA) or SuperSignal West Pico Plus Chemiluminescent Substrate (Thermo Scientific, #34578). A ChemiDoc XRS+ (Biorad) was used to visualize protein bands, which were subsequently quantified with ImageJ or ImageLab software (Biorad) and normalized to the corresponding to GAPDH, H3, or β-Actin bands.

### miRNA transfections

Wildtype and SATB1-KO SK-N-MC cells were plated on 6-well plates (Corning; 500,000 cells per well) and allowed to settle at 37°C with 5% CO_2_ for 24 hours. The following day, cells were transfected using the Lipofectamine RNAiMax Transfection Reagent (Thermo Scientific) with pre-warmed Opti-MEM serum-free media (Thermo Scientific). Scrambled miRNAs were used as a negative control. Wildtype cells were then transfected with a mimic of miR22-3p to see if this would lead to decreased GBA levels and SATB1-KO cells were transfected with an inhibitor of miR22-3p to see if this would lead to increased GBA levels (20 pmol miRNA per well). Media was changed 24 hours after transfection and 48 hours after transfection protein lysates were derived from cell cultures and Western blots were performed as described above.

### Chromatin Immunoprecipitation Sequencing (ChIP-seq)

A ChIP-seq approach was performed as previously reported (Riessland et al., 2019). In brief, we applied a modified protocol of the MAGnify Chromatin Immunoprecipitation System-Kit (Invitrogen) of ultra-sound-based sheared DNA and ChIP-sequencing libraries were generated using Ovation Ultralow System V2 Kit (NuGEN) according to the manufacturer’s instructions. To confirm quality and concentration of DNA libraries, we used both a Bioanalyzer (High Sensitivity DNA Chip (Agilent Technologies)) and the Tape Station (Agilent Technologies). Results from library sequencing, at the same molarities, were then multiplexed and sequenced on Illumina HiSeq 2500 sequencers using 100 single read and multiplexing conditions. In summary, all sequencing reads were tested for quality, trimmed, and aligned to the human genome. In order to crosslink the chromatin at day 60 of differentiation, cells were fixed for 8 minutes in fixation buffer containing 1% fresh formaldehyde (Thermo Scientific), washed in ice-cold PBS containing protease inhibitors (Roche), and subsequently sonicated (Covaris) using a chromatin shearing protocol in order to generate chromatin fragments of 100-400 bp in size. It is important to note that correct chromatin shearing has been successfully confirmed using the same DA neurons. After chromatin fragmentation, chromatin immunoprecipitation (ChIP) was performed using a well-characterized SATB1 antibody (Riessland et al., 2019) and an unspecific antibody (ctrl. IgG, LifeTechnologies). Sequencing reads were tested for quality using the FastQC online software (http://www.bioinformatics.bbsrc.ac.uk/projects/fastqc) and for further downstream analysis the sequences were trimmed using Trimmomatic (Bolger et al., 2014). For alignment to the human genome and model-based analysis of the results, we used a combined approach using Bowtie (Langmead et al., 2009), Presq (http://smithlabresearch.org/software/preseq) and MACS2 (Zhang et al., 2008). Finally, statistical significance was calculated, and data were analyzed. ChIP-sequencing data were visualized using the open-source software Integrative Genomics Viewer (Robinson et al., 2011). To analyze the regulatory region of MIR22HG, the reference H3K9ac ChIP-seq data (derived from human substantia nigra) was downloaded from encode.org.

### Immunocytochemistry

As reported previously (Riessland et al., 2019), for immunocytochemistry (ICC) cells were grown on sterile glass coverslips, fixed in 4% PFA in D-PBS, and incubated at room temperature for 30 minutes. Samples were permeabilized with 0.5% Triton X-100, blocked with 10% Normal Donkey Serum (NDS, Jackson ImmunoResearch #017-000-121), and then incubated overnight at 4°C with primary antibody (His-Tag (1:500), Invitrogen #Ma1-21315, Glucocerebroside (1:250), Glycobiotech #RAS_0010). The following day, samples were incubated with secondary antibody for 1 hour at room temperature. Secondary antibodies used were Goat anti-Rabbit IgG (H+L) Secondary Antibody, Alexa Fluor 594, Goat anti-Rabbit IgG (H+L) Secondary Antibody, Alexa Fluor 633, and Goat anti-Mouse IgG (H+L) Secondary Antibody, Alexa Fluor 568 (all purchased from Thermo Scientific). When used, ActinGreen™ 488 ReadyProbes™ Reagent (Thermo Scientific # R37110) was added along with secondary antibodies according to the manufacturer’s instructions. Cells were mounted on Superfrost Plus microscope slides (Thermo Scientific) with Prolong Gold Antifade reagent containing DAPI mounting medium (Invitrogen #P36931) for subsequent imaging.

### Fluorescent Dyes for Imaging Mitochondria and Lysosomes

Fluorescent imaging of mitochondria and lysosomes was performed as previously described (Riessland et al., 2019). For mitochondrial imaging, a 15-minute incubation at 37°C in a 500 mM solution of MitoTracker™ Red CMXRos (Thermo Scientific #M7512) in serum-free media was used. For lysosomal imaging, a 15-minute incubation at 37°C in a 1µM solution of LysoTracker™ Deep Red (Thermo Scientific #L12492) in serum-free media was used. In both cases, after staining all samples where fixed with 4% PFA in D-PBS and subsequently mounted to slides as described above.

### Fluorescence Microscopy

Fluorescence imaging was performed as previously reported (Riessland et al., 2019). In brief, fluorescence images were collected using a Zeiss LSM 710 confocal microscope and subsequent quantification and analysis of the images were performed in Fiji using minimal adjustment of contrast and brightness to ensure optimal and accurate representation of data. Quantification of Parkin puncta was performed as previously described (Ivatt et al., 2014), using an anti-Parkin antibody (rabbit polyclonal anti-Parkin, CST #2132). Corrected total cell fluorescence was calculated using Fiji as integrated density – (area of selected cell*mean fluorescence of background reading) (El-Sharkawey, 2016).

### Electron Microscopy

To perform transmission electron microscopy (TEM) as previously described (Riessland et al., 2019), cells were grown on ACLAR film. In short, cells were then fixed with 4% formaldehyde and 2% glutaraldehyde in 0.1M sodium cacodylate buffer (pH 7.4), and post-fixed with 2.5% glutaraldehyde and 0.25% tannic acid in 0.1M sodium cacodylate buffer for 15 minutes. They were then fixed with 2.5% glutaraldehyde in 0.1M sodium cacodylate buffer for 15 minutes, fixed with 1% osmium tetra-oxide in sodium cacodylate buffer for 30 min on ice, and subsequently washed three times in 0.1M sodium cacodylate buffer (pH 7.4) for 5 min and then rinsed three times with water. Cells were stained with 1% uranyl acetate for 30 min at room temperature followed by 12 hours on ice, dehydrated in increasing concentrations of ethanol; 50%, 70%, 90%, and 100% using Pelco Biowave Pro microwave automatic protocol (TedPella, Inc.), and then infiltrated with Epon812 resin, using an increasing concentration of resin in acetone; 50%, 70%, and 100% using Pelco Biowave Pro microwave automatic protocol (TedPella, Inc.), for 24 hours on a rotating rack with replacement of resin three times before polymerization for 48 hours at 60°C. Areas of interest were selected under the light microscope and trimmed for re-mounting and microtome sectioning where 70 nm sections were cut and collected on electron microscope grids, and samples were counter-stained using 1% uranyl acetate and Sato’s lead stain (Proc. XIth Int. Cong. on Electron Microscopy, Kyoto. 1986, pp. 2181-2182). The sections were then imaged with 1000-10000x magnification using 120kV operated Jeol 1400 plus TEM. Lipid vesicles were identified based on their structure and appearance (Tirinato et al., 2014) and counted in each cell (cell numbers: WT n=88, KO n=110). Statistical quantification was performed using GraphPad prism.

### Senescence-associated β-galactosidase (SA-β-gal) Assay

To test whether cells displayed phenotypes consistent with entering a state of cellular senescence, a senescence-associated β-galactosidase (SA-β-gal) assay (Cell Signaling Technology, Kit #9860) was performed according to the manufacturer’s protocol. In brief, after the respective treatment, cells were fixed and incubated over night with X-gal as substrate for the SA-β-galactosidase at 37°C. Subsequently, the substrate solution was removed, cells were overlayed with 70% glycerol (in PBS) and imaged with a microscope (Accu-Scope, EXI-410, Skye Color Camera). Images were analyzed and quantified using Fiji and statistical analysis was performed using Excel (Microsoft) and Prism (GraphPad).

### Immunoprecipitation

Immunoprecipitation of Parkin was performed using rabbit polyclonal anti-Parkin (CST #2132) and the Dynabeads™ Protein G Immunoprecipitation Kit (Thermo Scientific #1007D) according to the manufacturers protocol, with the slight modification of protein lysates being incubated overnight at 4°C with the antibody conjugated beads rather than at room temperature for 30 minutes.

### Tris Solubility Fractionation

To isolate Tris-soluble and insoluble protein fractions, cells were initially lysed in a 1% Triton-X 100 containing Tris buffered saline solution. The samples where then centrifuged and the supernatant was collected as the soluble fraction, and the Tris-insoluble pellet was re-suspended in 1X Laemmli buffer.

### GBA enzymatic activity assay

To measure the enzymatic activity of GCase (GBA), the GBA enzymatic activity assay (Novus bio) was used according to the manufacturer’s protocol as previously described (Riessland et al., 2019). In brief, this method utilizes p-nitrophenyl-β-D-glucopyranoside which is hydrolyzed specifically by GBA into a yellow-colored product. After incubation with the substrate, a colorimetric measurement was used to determine reaction rate which is directly proportional to enzymatic activity. Values were then normalized to protein content using a BCA assay as described above.

### Cathepsin D Enzymatic Activity Assay

As previously described (Riessland et al., 2019), Cathepsin D enzymatic activity was assessed using the Cathepsin D Activity Fluorometric Assay Kit (BioVision) according to the manufacturers protocol. Values were then normalized to protein content using a BCA assay as described above.

### Mice

Mice (C57BL/6) were housed in rooms on a 12 h dark/light cycle at 22 °C and their feeding was based on rodent diet (Picolab) and water available ad libitum. Mice were housed in groups of up to five animals except for mice that underwent stereotaxic surgery, which were housed singly to ensure recovery and avoid fighting. For all reported experiments, male mice were used. All animal experiments were approved by the Rockefeller University Institutional Animal Care and Use Committee and all described procedures were performed according to the guidelines described in the US National Institutes of Health Guide for the Care and Use of Laboratory Animals, and the ARRIVE guidelines.

### Cell survival

To assess cellular viability, the CCK8 cell viability kit (Dojindo) was used according to the manufacturer’s protocol in which the ability of cells to convert a water-soluble tetrazolium salt to a yellow-color formazan dye, which is soluble in the tissue culture media, is utilized and the measured amount of the dye, produced by dehydrogenases in the cells, represents cell viability and is directly proportional to the number of living cells. In brief, to determine viability, at the time of measurement cell medium was replaced by fresh medium containing 10% of CCK8 solution in serum-containing media. To determine the background, colorimetric absorbance was immediately measured at 450nm using the plate reader SpectraMax iD3 (Molecular Devices). Cells were incubated at 37°C for 2h hours and absorbance was measured again at 450nm.

### Viruses

As described previously (Riessland et al., 2019), the viruses used for stereotaxic injections were obtained from Vector Biolabs. The AAV1-EGFP-U6-shRNA virus was used as a control (scrambled shRNA and EGFP, #7040) and as previously reported, the shRNA virus for the silencing of *Satb1* was custom-made (Brichta et al., 2015). The *Satb1* shRNA construct used was designed to target bp 2329–2349 of *Satb1* mRNA (reference sequence: BC011132.1) and included the coding sequence for EGFP. GBA overexpression was achieved with the virus AAV1-hSYN1-mCherry-P2A-mGBA-WPRE. The control virus injected was AAV1-hSYN1-mCherry-WPRE. Viruses were packaged into a solution and injected at a concentration of 1 × 10^13^ genome copies per ml.

### Stereotaxic surgery

Stereotaxic injections were carried out as previously described (Brichta et al., 2015). In brief, we used an Angle Two stereotaxic frame for mouse with motorized nanoinjector (Leica) and injected 10-week-old male C57BL/6 WT mice (Charles River Laboratories) anesthetized with ketamine and xylazine. The experimental viruses (total injection volume 0.5 μl) were stereotactically injected targeting the ipsilateral SNpc (AP: −3.0 mm; ML: −1.2 mm; DV: −4.3 mm) and the control viruses were injected into the contralateral SNpc (AP: −3.0 mm; ML: +1.2 mm; DV: −4.3 mm). The injection rate was 0.05 μl/min using Hamilton syringes (30 gauge). Surgery wounds were sutured, and recovery was monitored after the injections.

### Histology

As previously described (Riessland et al., 2019), three weeks post injection mice were anesthetized with pentobarbital and transcardially perfused using PBS (pH 7.4), followed by 4% paraformaldehyde in PBS. Brains were postfixed in 4% paraformaldehyde in PBS at room temperature for 1 hour and then cryopreserved using a gradient of 5%, 15% and 30% sucrose. The brains were embedded in Neg-50 (Thermo Scientific), frozen and stored at −80 °C, and cut into 14-µm-thick coronal sections using a Microm cryostat and thaw-mounted onto Superfrost Plus microscope slides (Thermo Scientific). For staining of TH, brain sections were washed in PBS and permeabilized with 0.2% Triton X-100 in PBS, followed by blocking with 2% donkey serum and 0.1% fish gelatin in 0.2% Triton X-100 in PBS. Sections were then incubated with rabbit polyclonal anti-TH antibody (Millipore, #AB152) at a concentration of 1:250 overnight. The next day, slides were washed with PBS, incubated with Alexa Fluor 633 goat anti–rabbit IgG (Life Technologies, #A21070) for 2 hours at room temperature, and then after another washing step were mounted with ProLong Diamond Antifade containing DAPI (Life Technologies, #P36962), coverslipped, and stored in the dark until imaging.

### Blue Native Gel Electrophoresis and Western Blotting

As previously described (Riessland et al., 2019), for the assessment of mitochondrial complexes, cell lysates were prepared using the NativePAGE Sample Prep Kit (Life Technologies) and solubilized with 1% digitonin. A BCA assay (Thermo Scientific) was used to determine the protein concentrations as described above. Equal amounts of protein were then loaded onto a NativePAGE Novex 4-16% Bis-Tris Protein Gel (Life Technologies). A wet blotting method was used to transfer the proteins onto a PVDF membrane (Life Technologies, 0.45 µm), the membrane was then incubated with 8% acetic acid for 15min and washed with methanol and water before being blocked with 5% BSA in 20 mM Tris, 150 mM NaCl, and 0.1% (w/v) Tween 20, pH 7.5, for 2 hours. The membrane was subsequently immunoblotted with the respective primary antibody at 4 °C overnight. The primary antibodies used were NDUFA9 (a CI Subunit) (1/2500 by vol; ab14713: Abcam), UQCRC2 (a CIII Subunit) (1/2500 by vol; ab203832: Abcam), or MT-CO2 (a CIV Subunit) (1/2500 by vol; ab110258: Abcam). Primary antibodies were detected using either HRP-linked donkey anti–rabbit IgG (GE Healthcare, #NA934V; 1:10,000) or HRP-linked sheep anti– mouse IgG (GE Healthcare, #NA931V, 1:10,000) together with Western Lightning Plus-ECL (Perkin Elmer, #NEL105001EA).

### Measurements of cellular respiration

As previously described (Riessland et al., 2019), to measure cellular respiration as a reflection of cellular mitochondrial activity, we applied the XF mito stress kit using the Seahorse XFe96 Analyzer (Agilent) according to the manufacturer’s protocol. In brief, we plated out 40,000 cells per well on a 96-well plate one day prior the measurement and incubated cells in 5% CO_2_ at 37°C overnight. On the day of measurement, cells were washed with XF cell mito stress test assay medium (Agilent) and incubated in the medium for one hour prior to the measurement in a CO_2_-free incubator at 37°C. During the measurement program, cells were challenged with 1.0 µM Oligomycin (Port A), 0.5 µM FCCP (Port B) and 0.5 µM Rotenone/antimycin A (Port C). Results of the measurement were subsequently analyzed using the Wave software (Agilent). In order to measure mitochondrial activity in hESC-derived DA neurons, approximately 40,000 neurons which express NURR1::GFP were plated out at day 25 of differentiation into DA neurons. The cells were then treated as described above, with the exception of 1.5 µM FCCP (Port B) being used during measurement. After the Seahorse run, results were normalized on the protein concentration of every individual well (based on BCA assay).

### hESC culture and differentiation

The hESCs wild-type H9 (WA-09), SATB1^KO^, NURR1-GFP reporter line were maintained using E8-essential medium (Fisher Scientific) without feeder on VTA-N (Fisher Scientific) and passaged every 4-5 days by EDTA. Midbrain dopamine (DA) differentiation from hESC was done with a protocol previously published by our group (Kim et al., 2021; Riessland et al., 2019).

### ChIP-seq, ATAC-seq and RNA-seq of wildtype and SATB1-KO DA neurons

ChIP-seq, ATAC-seq and RNA-seq was described and performed previously (Riessland et al., 2019) and data is available at: https://www.ebi.ac.uk/biostudies/arrayexpress/studies/E-MTAB-5965/.

### 2-deoxyglucose experiment

N2A cells were treated for 48h with the glycolysis inhibitor 2-deoxyglucose (500nM, Sigma). Cells that rely on glycolysis cannot survive with 2-deoxyglucose. Cell survival was determined using a CCK8 assay (Dojindo) according to the manufacturers protocol as described above.

### Cell treatments with CCP, 6-OHDA or H2O2

6-OHDA was always freshly prepared in 0.9% NaCl+0.05% ascorbic acid and used immediately to prevent oxidation. When 6-OHDA was used, the mock treatment was 0.9% NaCl+0.05% ascorbic acid. Water served as mock treatment for H2O2. Cells were treated with DMSO (Sigma) or 2 uM CCCP (Sigma) for 3.5 hours, or CCCP followed by a 0.5-hour washout with serum free media.

### Dot blots for GluCer

Whole cell lysates (in RIPA buffer) were used for dot blots. The dots were normalized on protein concentration (BCA assay) and cell numbers (EVE Automated Cell Counter, VWR). Each drop contained 10ug protein in 5 ul RIPA. Drops were placed on nitrocellulose membranes. After a short drying period (20 minutes), membrane was visualized and imaged with Ponceau S. Staining Solution (Thermo Scientific) for standardization and then incubated in TBS-T + 5% BSA for 1h on RT. After this blocking step, membrane was incubated with anti-glucocerbroside antibody overnight at 4°C in TBS-T + 5% BSA. Then membrane was washed 5 times for 5 minutes in TBS-T. Subsequently, membrane was incubated with secondary antibody labeled with HRP for 1h in TBS-T + 5% BSA. After 5 additional washes (5 minutes each), Western Lightning Plus-ECL (Perkin Elmer, #NEL105001EA) was used to expose western blot detection films which were developed in the dark room to visualize lipid dots, which were subsequently quantified with ImageJ or ImageLab (Biorad).

### Plasmid transfection

The following plasmids have been used in the manuscript: human SATB1 (Origene), human GBA (Origene), human a-SYN (A53T mutant) (Addgene), and mt-Keima (Addgene). The alphasynuclein-A53T plasmid was a gift from David Rubinsztein (Addgene plasmid # 40823; http://n2t.net/addgene:40823; RRID:Addgene_40823) (Furlong et al., 2000); mt-mKeima was a gift from Richard Youle (Addgene plasmid # 131626; http://n2t.net/addgene:131626; RRID:Addgene_131626) (Vargas et al., 2019). mKeima-Red-Mito-7 was a gift from Michael Davidson (Addgene plasmid # 56018; http://n2t.net/addgene:56018; RRID:Addgene_56018) Transfections have been performed with Lipofectamin 3000 (Thermofisher) according to the manufacturer’s protocol.

### SK-N-MC and Neuro-2A cells

SK-N-MC cells (ATCC) and Neuro-2A (N2A) cells (ATCC) were cultured according to the vendor’s protocol. In brief, cells were grown at 37C in 5% CO2. The used medium for both cell lines is ATCC-formulated Eagle’s Minimum Essential Medium (Catalog No. 30-2003). 1% Pen/Strep and fetal bovine serum to a final concentration of 10% was added.

### rhGCase experiments

For the enzyme replacement experiments, cells were treated for 24h with 100ng/ml media of human recombinant GBA (rhGBA-His (R&D Systems)).

### GluCer treatment in N2A cells for SA-β-gal

Senescence-associated β galactosidase assay was performed according to the manufacturer’s protocol and as described above (Cell Signaling Technology, Kit #9860). 5,000 cells were plated on each well of a 96-well dish. 24 hours later medium containing DMSO, 2.5, 5, 10, 20, and 40uM Glucocerebrosides (Avanti, USA) was added. The β-gal assay was performed 7-days after treatment was added and Fiji was used for quantification of senescent cells as described above.

### GluCer treatment in mDA neurons for SA-β-gal (TR)

Mature WA09 DA neurons were treated for 7 days with 2.5 uM GluCer, 40 uM GluCer, or DMSO control in normal culture media (replenished every other day with media changes) prior to SA-β-gal assay, as described above.

### GluCer treatment in N2A cells for lysotracker

Cells were treated with 2.5 uM GluCer, 40 uM GluCer, or DMSO control in normal culture media for 6 days prior to staining with Lysotracker Deep Red, as described above.

### RNA isolation and RNAseq of GluCer-treated DA neurons

RNA was isolated from mature wildtype DA neurons following 7-day treatment with GluCer using the RNeasy Micro Kit (QIAGEN) and sent for bulk RNAseq to Azenta. In brief, sample quality control and determination of concentration was performed using TapeStation Analysis by Azenta, followed by library preparation and sequencing. Computational analysis included in their standard data analysis package was used for data interpretation.

