## Supplementary Figures for "The SATB1-MIR22-GBA axis mediates glucocerebroside accumulation inducing a cellular senescence-like phenotype in dopaminergic neurons"

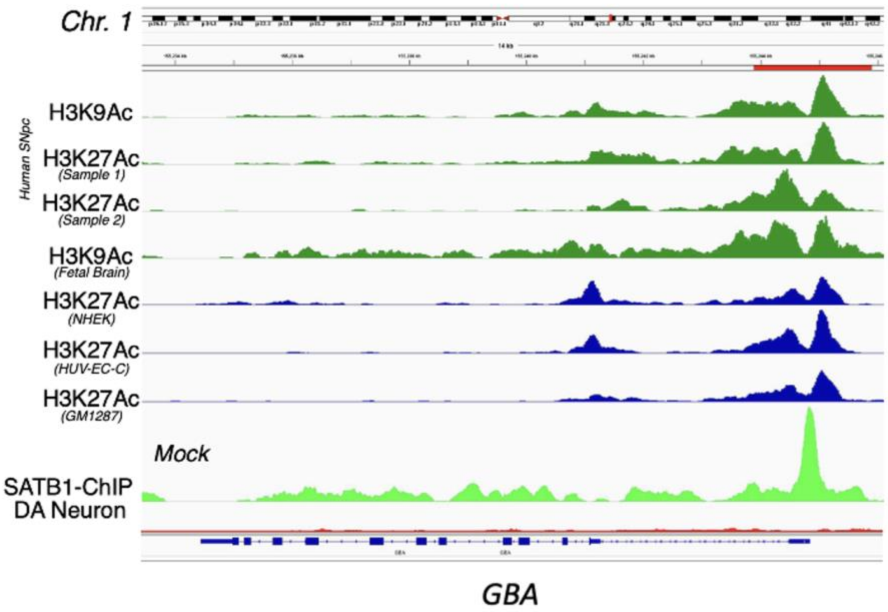


**Figure S1. SATB1 binds to the regulatory region of GBA in dopaminergic (DA) neurons.** ChIP-seq analysis reveals significant binding of SATB1 to the GBA gene in 60-day-old human embryonic stem cell–derived DA neurons. The GBA gene is overlaid with H3K27Ac and H3K9Ac enrichment tracks from human substantia nigra samples, human fetal brain, and three human cell lines. H3K27Ac and H3K9Ac data were obtained from http://www.roadmapepigenomics.com. The chromosomal locus and exons of GBA are shown, with the regulatory region of GBA highlighted by a red line. ChIP-seq was performed in quadruplicate (n=4).

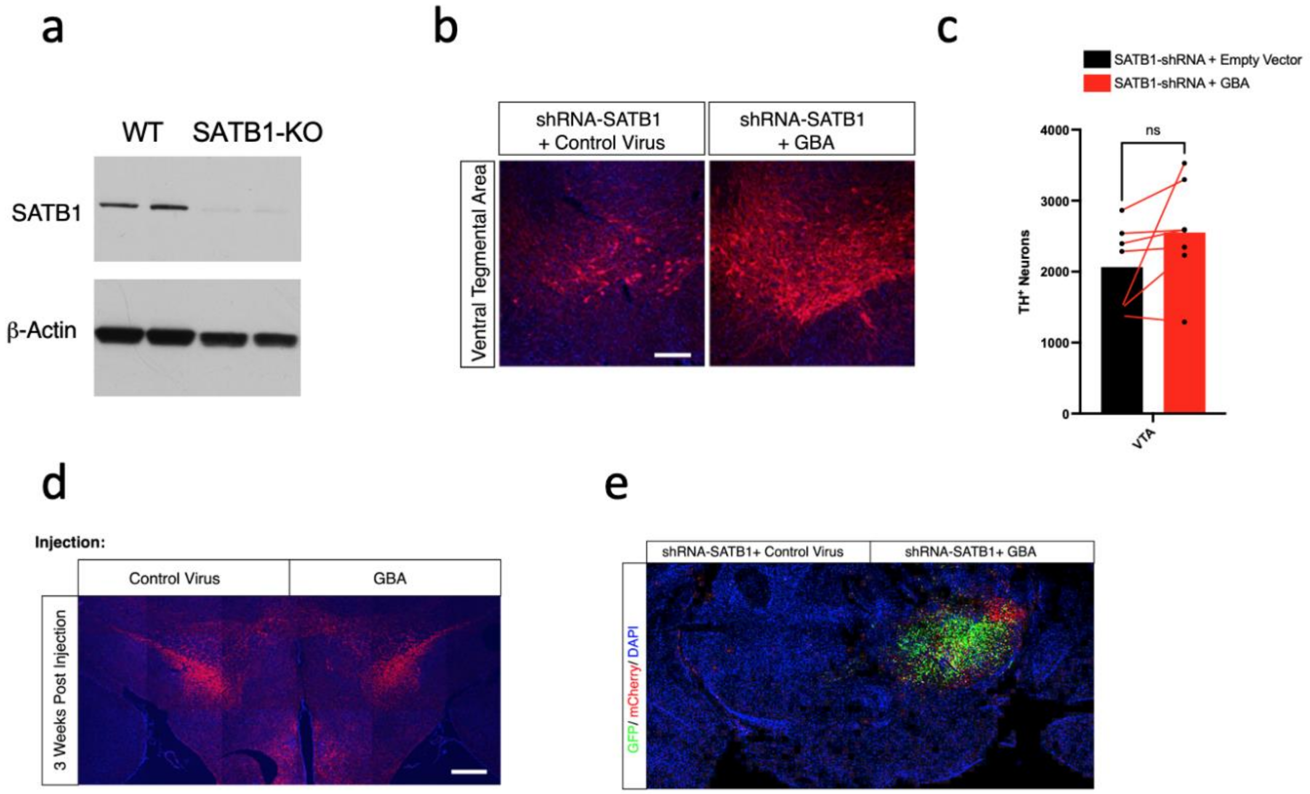


**Figure S2. Confirmation of SATB1 knockout (KO) and *in vivo* virus experiments a,** Western blot analysis confirms the elimination of SATB1 protein in N2A SATB1-KO cells. **b** and **c,** Representative images and quantification of tyrosine hydroxylase (TH) immunofluorescent staining in mice that received a stereotaxic injection with a shRNA-SATB1 virus and a control vector or a contralateral injection with a shRNA-SATB1 virus and a GBA-overexpressing virus. The ventral tegmental area (VTA) was analyzed using unbiased stereological cell counting of $TH^+$ cells, showing no significant effect (n=7). Data are presented as mean ± S.E.M. Student's t-tests were performed. ** p<0.01; *** p<0.001. **d** and **e,** Viral overexpression of GBA in mouse dopaminergic neurons did not affect TH expression and survival of $TH^+$ neurons compared to the opposite midbrain injected with the empty vector control virus, observed 3 weeks following viral injection. Scale bar: 500 mm.

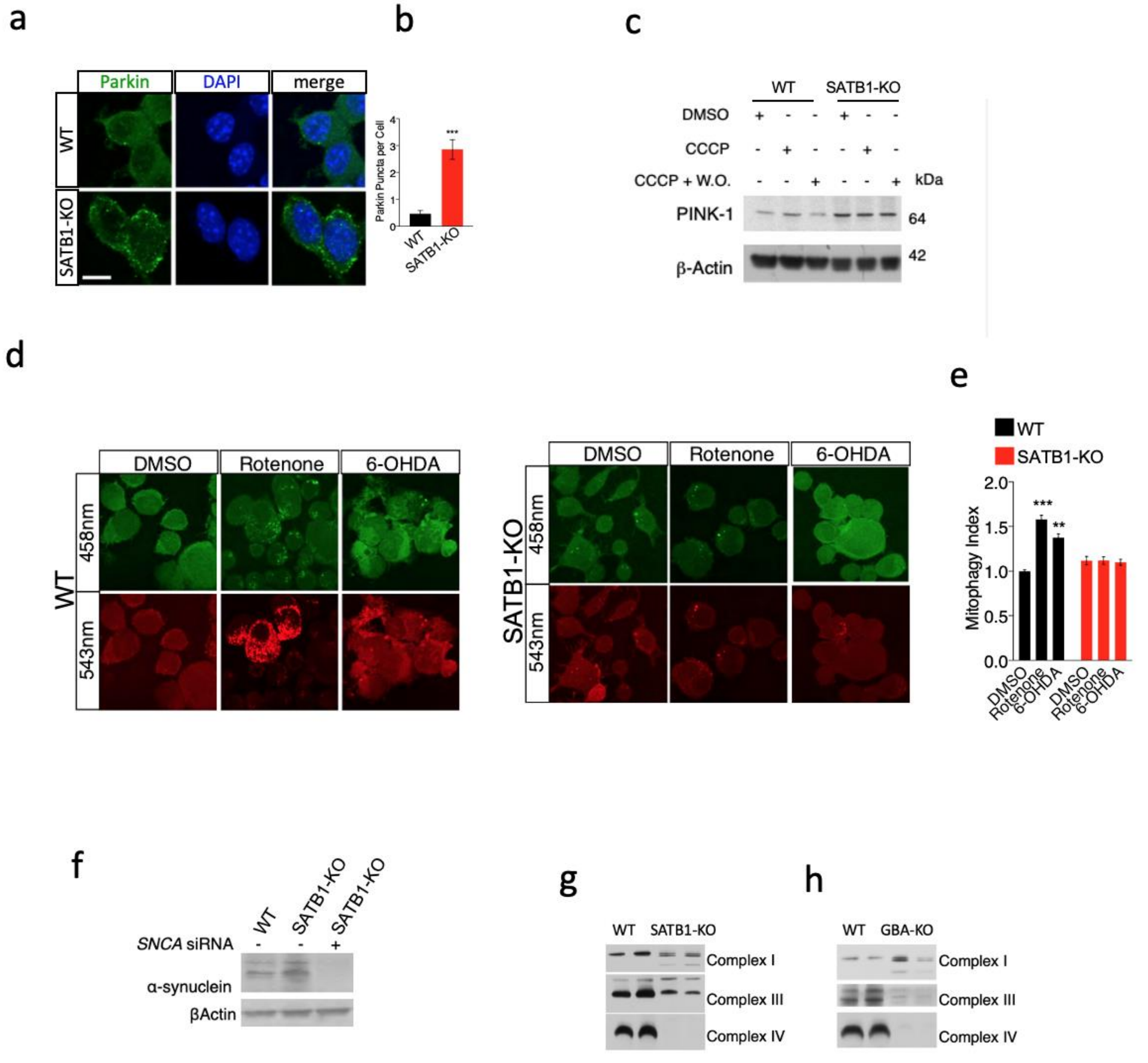


**Figure S4. SATB1 knockout (KO) leads to accumulation of dysfunctional mitochondria and altered mitophagy index.**

**a, b**, Immunofluorescent staining using anti-Parkin antibodies revealed a significant increase in Parkin-labeled puncta in N2A$^{SATB1\text{-}KO}$ cells compared to controls. **c**, Immunoblotting analysis of PINK1 protein levels after treatment with DMSO or CCCP for 3.5 hours or CCCP followed by a 0.5-hour washout with serum-free media. PINK1 protein levels increased with CCCP treatment and returned to baseline after washout in wild-type (WT) N2A cells. However, N2A$^{SATB1\text{-}KO}$ cells showed elevated basal levels of PINK1 protein, which remained unchanged following treatment (n=4). **d, e**, Functional analysis of mitophagy in WT and SATB1-KO N2A cells transfected with mt-mKeima plasmid and treated with rotenone or 6-OHDA. Total mitochondrial fluorescence (ex.: 458 nm) is shown in green, and mitophagy (ex.: 543) is shown in red. The mt-mKeima quantification was performed with at least 100 cells analyzed per condition. The 'mitophagy index' was calculated as the ratio of fluorescence intensity emitted from the two excitation peaks: 543 nm divided by 458 nm, as previously described (Goiran et al., 2022). Data are presented as mean ± S.E.M. Student's t-tests were performed. * $p<0.05$; ** $p<0.01$; *** $p<0.001$. **f,** Western blot confirmation of α-SYN knockdown in N2A cells. **g, h** Native gel electrophoresis showing loss of oxidative phosphorylation complex integrity in both SATB1-KO and GBA-KO cells.

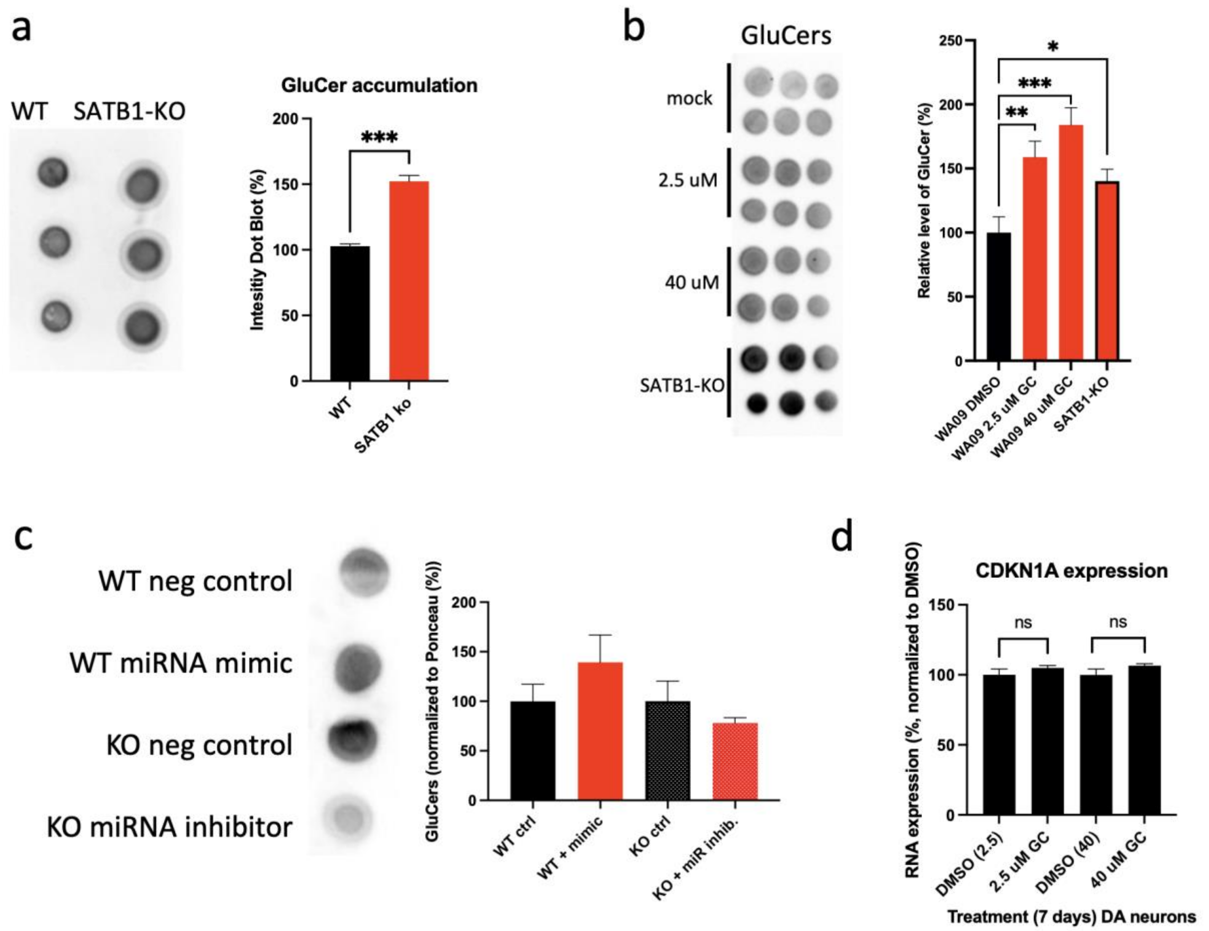


**Figure S5. Dot blot analysis of GluCer in the N2A cell line, lipid-treated dopaminergic (DA) neurons, and the miRNA-treated SK-N-MC cell line.**

**a**, Dot blot analysis of GluCer in the N2A cell line showing increased expression in SATB1-KO DA neurons. **b,** Dot blot analysis of GluCer in mature wild-type (WT; WA09) human DA neurons treated with DMSO control, 2.5 uM GluCer, or 40 uM GluCer, and SATB1-KO DA neurons showing similar lipid accumulation in lipid-treated and SATB1-KO DA neurons. **c,** Dot blot analysis of GluCer in WT SK-N-MC cells treated with a miR22-3p mimic and SATB1-KO SK-N-MC cells treated with a miR22-3p inhibitor showing a non-significant but trending increase in GluCer with the addition of the miR22-3p mimic and a decrease in GluCer with the addition of the miR22-3p inhibitor. **d**, Quantification of *CDKN1A* (p21) in WT DA neurons did not show any significant changes.
